# scGHOST: Identifying single-cell 3D genome subcompartments

**DOI:** 10.1101/2023.05.24.542032

**Authors:** Kyle Xiong, Ruochi Zhang, Jian Ma

**Affiliations:** Computational Biology Department, School of Computer Science, Carnegie Mellon University, Pittsburgh, PA 15213, USA

## Abstract

New single-cell Hi-C (scHi-C) technologies enable probing of the genome-wide cell-to-cell variability in 3D genome organization from individual cells. Several computational methods have been developed to reveal single-cell 3D genome features based on scHi-C data, including A/B compartments, topologically-associating domains, and chromatin loops. However, no scHi-C analysis method currently exists for annotating single-cell subcompartments, which are crucial for providing a more refined view of large-scale chromosome spatial localization in single cells. Here, we present SCGHOST, a single-cell subcompartment annotation method based on graph embedding with constrained random walk sampling. Applications of SCGHOST to scHi-C data and single-cell 3D genome imaging data demonstrate the reliable identification of single-cell subcompartments and offer new insights into cell-to-cell variability of nuclear subcompartments. Using scHi-C data from the human prefrontal cortex, SCGHOST identifies cell type-specific subcompartments that are strongly connected to cell type-specific gene expression, suggesting the functional implications of single-cell subcompartments. Overall, SCGHOST is an effective new method for single-cell 3D genome subcompartment annotation based on scHi-C data for a broad range of biological contexts.

## Introduction

The development of high-throughput three-dimensional (3D) whole-genome mapping methods, such as Hi-C [1], has facilitated the delineation of multiscale 3D genome features, including A/B compartments [1], subcompartments [2, 3], topologically associating domains (TADs) [4, 5], and chromatin loops [2]. Numerous studies have demonstrated that these 3D genome features are intertwined with important genome functions, such as gene transcription and DNA replication [6, 7]. A major challenge in studying 3D genome structure and function is revealing the 3D genome features and their variability at single-cell resolution [8]. Emerging single-cell 3D genome mapping technologies, particularly single-cell Hi-C (scHi-C), have enabled the analysis of higher-order chromatin structure in individual cells [9–15]. Recently, several new computational methods have been developed to address the analysis challenges posed by the high data sparsity of scHi-C, enhancing overall data quality [16–18] and characterizing multiscale 3D genome features and their heterogeneity at single-cell resolution [17, 19, 20].

Although computational methods exist for calling A/B compartments [13, 17], TADs [17], and chromatin loops [20] from scHi-C data, no method currently reveals single-cell 3D genome subcompartments. The 3D genome exhibits critical subcompartment patterns that the binary A/B compartment definitions do not adequately capture [21]. For bulk Hi-C data, Rao et al. [2] first refined the A/B compartments into five major subcompartments (A1, A2, B1, B2, and B3) in GM12878 cells using high-coverage interchromosomal Hi-C contact maps, which show distinct correlations with various epigenomic features. More recent methods have been developed to identify subcompartments based on low-coverage bulk Hi-C in different cell types, including Calder [22], SCI [23], and our own earlier work, SNIPER [3]. Collectively, these studies show that subcompartment annotations significantly advance the binary A/B compartment definitions with strong stratification of functional genomic signals, providing key insights into genome structure-function connections. However, applying current subcompartment annotation methods developed for bulk Hi-C to scHi-C data remains infeasible primarily due to two challenges: (1) scHi-C data requires specialized methods to mitigate its high data sparsity and noise for identifying specific 3D genome features, such as subcomparments; (2) nearly all scHi-C datasets lack sufficient coverage to reveal interchromosomal chromatin interactions for facilitating subcompartment annotations in single cells, necessitating new approaches. Consequently, no method currently addresses these challenges for annotating 3D genome subcompartments based on scHi-C data.

Here, we develop a new method named SCGHOST (single-cell graph-based Hi-C organization and segmentation toolkit) that provides genome-wide annotation of subcompartments in each individual cell based on scHi-C data. Our method leverages the data imputed from our recently developed algorithm, Higashi [17]. Importantly, SCGHOST employs graph-embedding neural networks trained on a unique, constrained random walk sampling strategy to partition scHi-C contact maps into subcompartment annotations. By applying SCGHOST to scHi-C data in several cell lines as well as single-cell 3D genome imaging data, we find that SCGHOST is capable of revealing single-cell subcompartments that offer further insights into the functional implications of chromatin spatial localization in individual cells. Moreover, SCGHOST uncovers novel cell type-specific connections between subcompartments and gene transcription based on scHi-C data from human prefrontal cortex. Together, SCGHOST is expected to be a highly useful new method for enhancing the scHi-C analysis toolbox for studying single-cell 3D genomes.

## Results

### Overall design of the SCGHOST framework

SCGHOST is a single-cell compartmentalization framework and views scHi-C contact maps as graphs, where genomic loci are vertices in the graph and are connected through edge weights defined by Hi-C contact frequencies among loci. SCGHOST employs a unique random sampling procedure that filters noise in imputed scHi-C data, represents (embeds) each genomic locus (graph vertex) in single cells as a continuous-valued vector, and uses unsupervised learning to discretize single-cell genomes and identify 3D genome subcompartments (see **Fig**. 1a for an overview).

**Figure 1:**
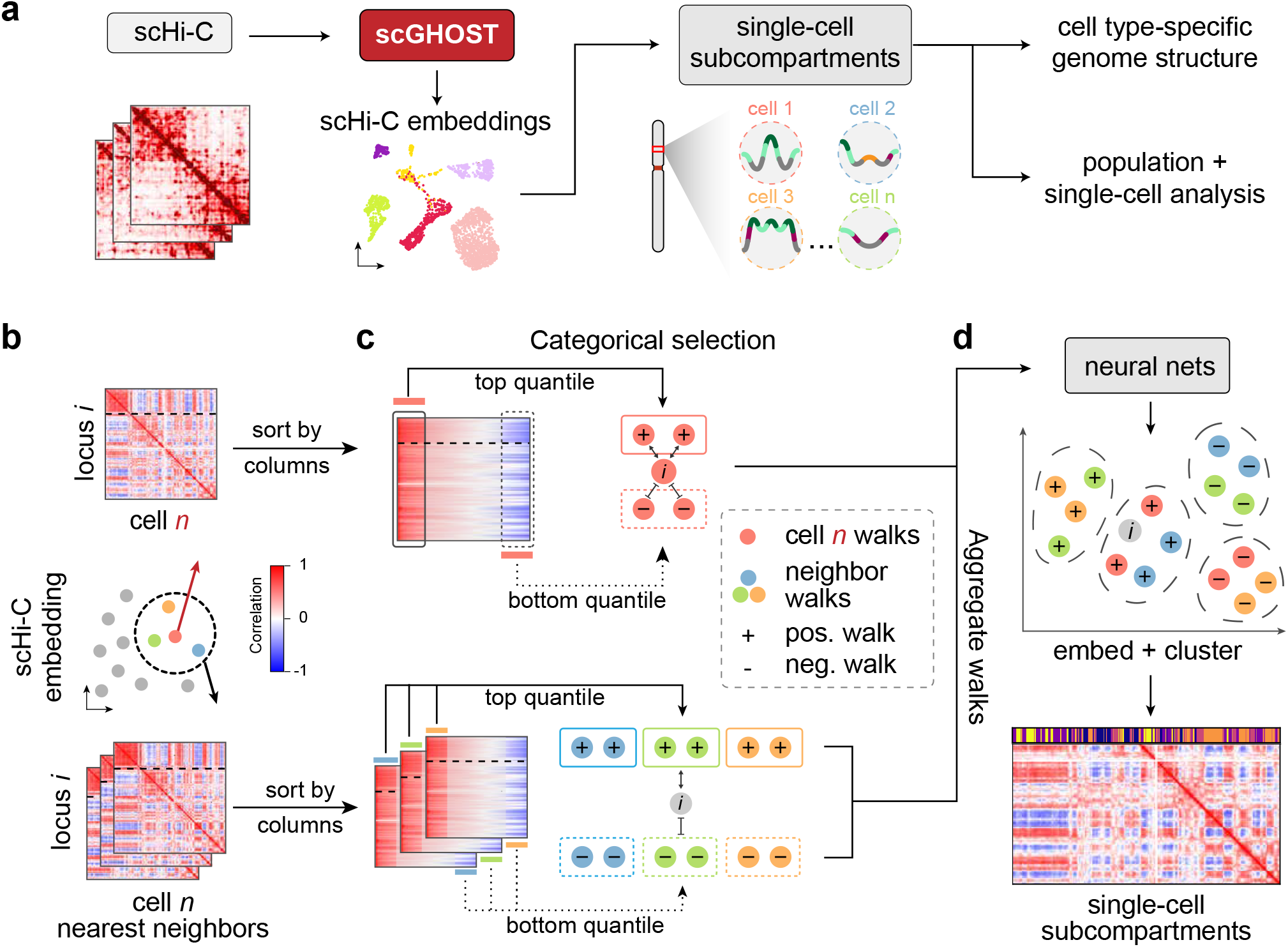
Overview of the SCGHOST framework. **a**. Schematic of the SCGHOST workflow. **b**. Higashi embeddings are used to identify the cells that exhibit the most similarity in the single-cell embedding space. **c**. Random walks create sparse graphs that portray the most crucial connections among genomic loci. **d**. Walks are aggregated and fed into a graph embedding model, which generates embeddings of each genomic locus. These embeddings are subsequently clustered and compared to reach a final set of subcompartment annotations comparable across chromosomes and cells.

The input of SCGHOST consists of the imputed scHi-C contact maps of a cell and its nearest neighbors, defined by the Euclidean distance between the scHi-C embeddings of the cell and all other cells in the scHi-C dataset. Our framework then outputs a set of discrete annotations for each genomic locus in each individual cell. These annotations can reveal cell-to-cell variability of subcompartments and facilitate the analysis of cell type-specific genome structure and function.

SCGHOST comprises four main components. (1) A new sampling procedure that estimates the most reliable genomic interactions in a single cell using imputed scHi-C contact maps and the maps of cells whose Higashi scHi-C embeddings have the lowest Euclidean distance from it **(Fig**. 1b). This step outputs a sparse, undirected, weighted graph containing only the strongest Hi-C contacts among genomic loci **(Fig**. 1c). (2) A graph embedding procedure that embeds each genomic locus in scHi-C maps and aggregates random walks across multiple cells, connecting loci that should have contacts but otherwise would not due to noise in scHi-C data **(Fig**. 1d). The embeddings are used in the subsequent clustering step for annotating single-cell subcompartments. (3) A clustering process unique to scHi-C data that ensures clusters in different chromosomes correspond to the same set of genome-wide subcompartments. (4) A procedure that makes single-cell subcompartment annotations comparable in all single cells **(Fig**. 1d). Overall, SCGHOST effectively facilitates the discovery of single-cell subcompartments by cohesively segmenting genomes of individual cells into discrete spatial states using scHi-C data.

### SCGHOST identifies distinct subcompartments from scHi-C data of GM12878 cells

We first applied SCGHOST to the Higashi-imputed scHi-C data of GM12878 at 500kb resolution [12]. In **Fig**. 2a, we compare the median intra- and inter-subcompartment observed-over-expected contact frequencies across all cells. We found that regions in all single-cell subcompartments, denoted scA1, scA2, scB1, scB2, and scB3, preferentially interact with regions within the same subcompartment. Intra-cluster interactions are significantly more frequent than inter-cluster interactions, with a one-sided p-value of less than 0.001 (see **Supplementary Note**). These interaction patterns among single-cell subcompartments suggest that SCGHOST possesses the sensitivity to categorize scHi-C contact maps into distinct subcompartments with unique Hi-C contact patterns.

**Figure 2:**
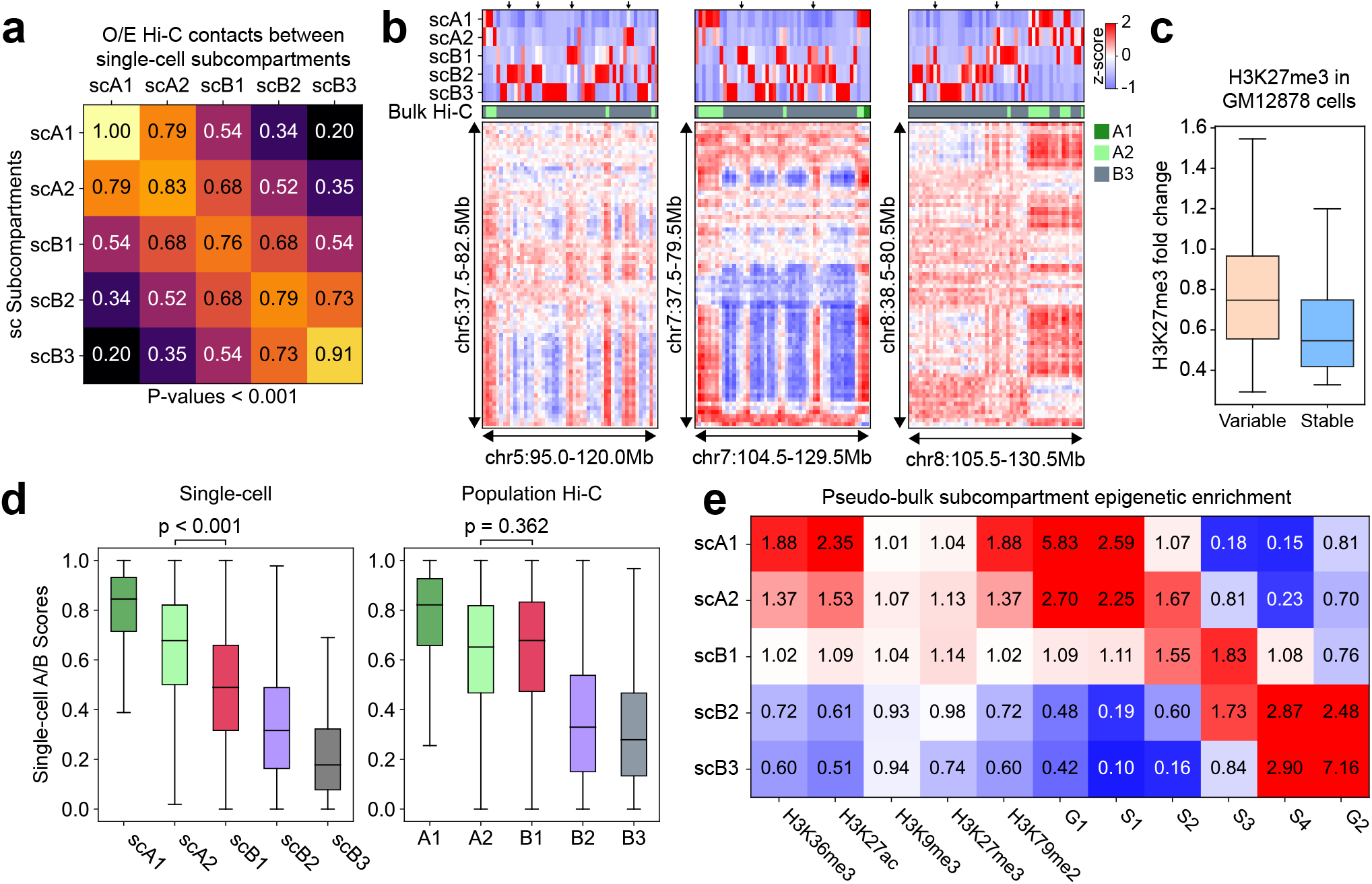
SCGHOST’s application to GM12878 single-cell Hi-C data showcases its accuracy in annotating single-cell subcompartments. **a**. Median observed-over-expected contact frequencies among single-cell subcompartments. **b**. Example regions, indicated by the arrows at the top, where single-cell subcompartments of the population of GM12878 cells correlate more strongly than bulk Hi-C subcompartments with contact patterns in the pseudo-bulk Hi-C matrix. **c**. The distribution of H3K27me3 in GM12878 genomic regions with variable and stable single-cell subcompartment annotations (*p <* 0.001). **d**. Boxplots displaying the distribution of Higashi single-cell A/B scores in single-cell subcompartments (left) and population subcompartments (right). **e**. The epigenomic signal enrichment of pseudo-bulk subcompartments computed from single-cell subcompartments.

Next, we sought to demonstrate that SCGHOST subcompartments could enhance the overall genome segmentation compared to bulk Hi-C subcompartments. We found that the aggregated single-cell subcompartment annotations correspond more closely to changes in scHi-C interaction patterns than bulk Hi-C annotations (**Fig**. 2b). In Fig. 2b, we show an example at the population level where fluctuations in contact patterns in the B3 subcompartment more closely resemble A2 (see indicated regions in Fig. 2b). In annotations from bulk Hi-C, these regions remained annotated as B3, while SCGHOST placed them in more refined, active subcompartments. We also observed this on a genome-wide level **(Fig**. S1), where the aggregated single-cell subcompartments correspond to reduced variance of scHi-C contact frequencies. Therefore, at the population level, single-cell annotations from SCGHOST improve the segmentation from the original bulk Hi-C annotations [2], despite being distributed differently (**Fig**. S2).

We then utilized single-cell subcompartment annotations to better understand the single-cell variability of facultative heterochromatin, largely annotated as B1 subcompartments from bulk Hi-C and interacting with genomic loci in both the A and B compartments [2]. We partitioned genomic loci of GM12878 cells into variable and stable single-cell subcompartments (**Supplementary Note**) and calculated the enrichment of H3K27me3, the marker of facultative heterochromatin. In **Fig**. 2c, we observed significantly higher H3K27me3 enrichment over control in variable genomic regions than stable ones in the cell population (p <0.001). This suggests that facultative heterochromatin more likely consists of variable regions annotated as B1 from bulk Hi-C. Indeed, compared to stable subcompartments variable subcompartments are more highly associated with the B1 subcompartment (**Fig**. S3), which also exhibits variable single-cell annotations (**Fig**. S2). These results suggest that facultative heterochromatin regions are characterized by high cell-to-cell variability in spatial position, further confirming the reliability of the SCGHOST results.

While we have demonstrated that SCGHOST can improve single-cell genome segmentation compared to bulk Hi-C subcompartments, it is necessary to reconcile the discrepancy between single-cell subcompartments and bulk Hi-C subcompartments (**Fig**. S2). We compared single-cell A and B (denoted scAB) compartment scores corresponding to single-cell and bulk Hi-C subcompartments (**Fig**. 2d). A and B compartment scores for each cell were computed using Higashi-imputed scHi-C maps **(Supplementary Note**). In single-cell subcompartments, the distribution of scAB in all single-cell subcompartments are significantly distinct from one another (p <0.001). By contrast, scAB are not differently distributed by any significant margin in the A2 and B2 subcompartments from bulk Hi-C (p =0.362). *P*-values were calculated for χ^2^ = 1200 due to floating point accuracy limits of the t-test for greater χ^2^ values. Despite the differences between single-cell and bulk Hi-C subcompartment annotations, single-cell annotations from SCGHOST offer greater power for segmenting single-cell genomes with subcompartment patterns.

We found that single-cell subcompartment annotations overall consistently stratify epigenomic signals such as histone modifications and replication timing. We computed pseudo-bulk subcompartments using our GM12878 single-cell annotations (**Supplementary Note**) and constructed an epigenomic mark enrichment profile of our pseudo-bulk subcompartments (**Fig**. 2e). The trend of histone mark enrichment and early-late replication timing when transitioning from scA1 to scB3 is consistent with that in bulk HiC subcompartments [2, 3]. When aggregated in a population of single cells, SCGHOST subcompartments align with active/repressive histone marks and early-late replication timing.

Taken together, these results collectively demonstrate that SCGHOST can reliably and effectively identify single-cell subcompartment patterns using scHi-C data.

### SCGHOST unveils single-cell subcompartment associated with transcriptional variability

We next evaluated single-cell subcompartment annotations in the WTC11 human iPS cells [24] at 500kb resolution. Specifically, we utilized SCGHOST to uncover spatially variable and stable genomic loci in terms of subcompartment states in individual cells. For each genomic locus, we calculated the information content of subcompartment annotations across all cells from the expected distribution of subcompartments (**Supplementary Note**). Regions with high information content share the same subcompartment annotation across the majority of cells and are thus considered stable, while regions with low information content tend to display a more uniform subcompartment distribution (i.e., more variable).

For each genomic locus of WTC11 cells in the variable and stable states, we computed the fraction of cells in each subcompartment (**Fig**. 3a, blue). We found that the stable subcompartment regions are primarily annotated as scA1 and scB3, suggesting that the spatial positions of scA1 and scB3 regions are mostly conserved among cells. By contrast, genomic regions with variable spatial positions across cells are marked by relative uniform distributions of single-cell subcompartments **(Fig**. 3a, orange).

**Figure 3:**
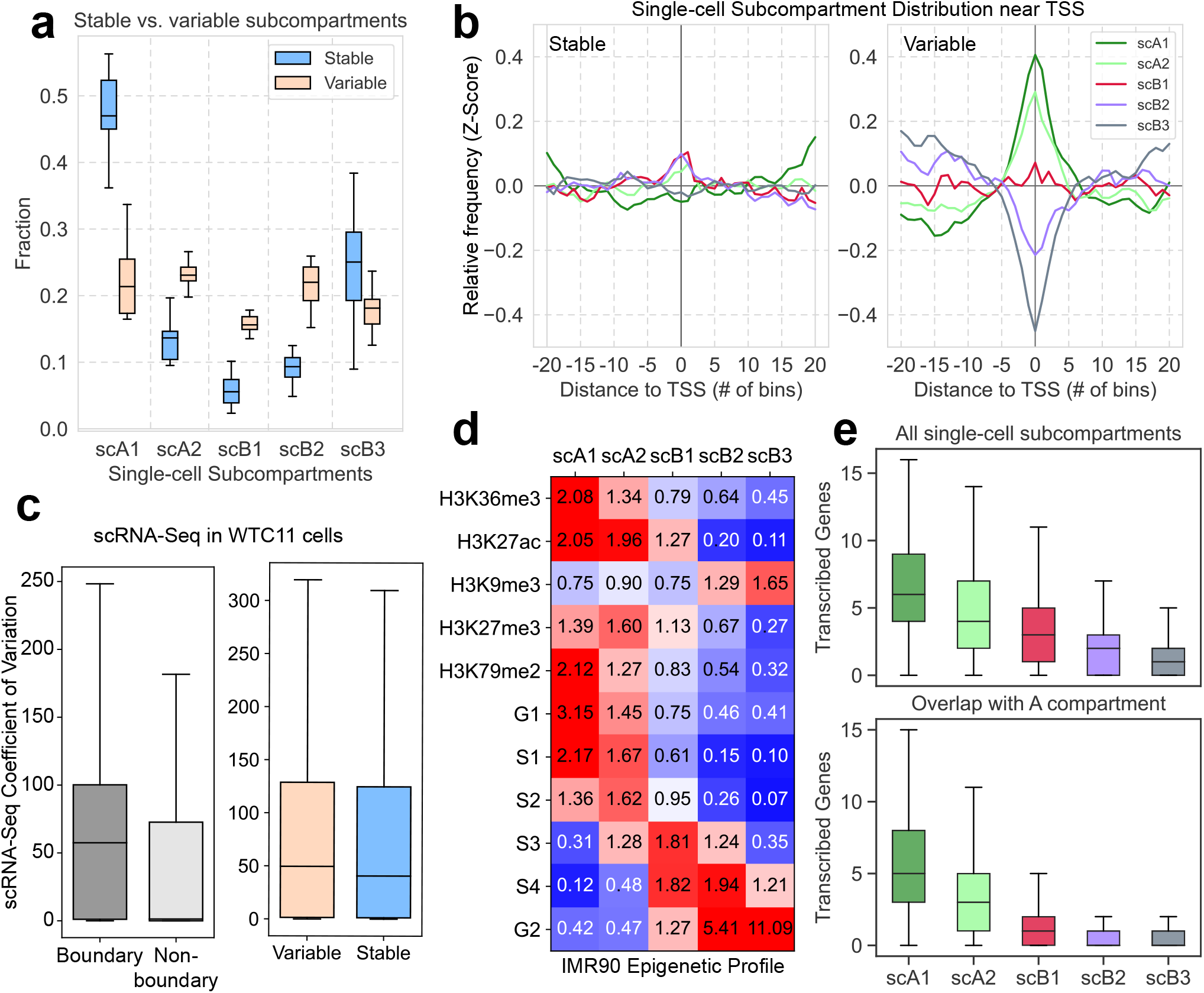
SCGHOST’s application to WTC11 scHi-C data and IMR90 single-cell 3D genome imaging data. **a**. Proportion of single-cell subcompartments in stable (blue) and variable (orange) chromatin states across all chromosomes. **b**. Relative frequency of single-cell subcompartments in 10 Mb regions flanking transcription start sites in stably (left) and variably (right) transcribed genes (defined based on the coefficient of variance of the MAGIC imputed WTC11 scRNA-seq.). **c**. scRNA-Seq expression is more variable in single-cell subcompartment boundary and variable subcompartment annotation regions. **d**. Epigenomic enrichment of pseudobulk subcompartments in chr21, as revealed by the IMR90 3D genome imaging dataset. **e**. Co-assayed gene transcription activity in single-cell subcompartments genome-wide (top) and in regions that overlap with A compartment annotated by bulk Hi-C (bottom).

We next examined the connection between the variability of subcompartments and gene transcription activities. We first categorized genes into stably and variably transcribed genes based on scRNA-seq. Specifically, we quantified the transcription variability as the coefficient of variance of the MAGIC [25] imputed scRNA-seq of WTC11 [26], which was further binarized into genes with stable and variable transcription activity based on quantiles. Near the transcription start sites (TSS) of stably transcribed genes, subcompartments are mostly uniformly distributed **(Fig**. 3b, left), whereas variably transcribed genes are more frequently located in scA1 and scA2 and less frequently in scB2 and scB3 **(Fig**. 3b, right).

When we divided these subsets of stably and variably transcribed genes by stable and variable subcompartment annotations, we found that stably transcribed genes in stable subcompartments are mostly marked by scB3 and in variable subcompartments are marked by scA2, scB1, and scB2 **(Fig**. S4a,b). However, variably transcribed genes in stable subcompartments are highly associated with scA1 and in variable subcompartments are associated with scA2 and scB1 (**Fig**. S4c,d). Notably, variably transcribed genes are consistently in the most active scA1 subcompartment. Furthermore, transcription of stable genes does not occur more frequently in the scA1 subcompartment. Together, stably transcribed genes are surprisingly marked by every subcompartment except scA1, whereas variably transcribed genes are primarily marked by scA1 in stable subcompartments, and scA2 and scB1 in variable subcompartments (**Fig**. S4).

We then aimed to show the connection between the variability of subcomparmentalization and transcription activity across the cell population. We used the aforementioned information content to define genomic regions with stable and variable subcompartment annotations. We observed that regions undergoing changes of subcompartmentalization across cells are more enriched of genes with more variable transcription activity (one-sided t-test p = 6.90 *×* 10^*−*3^; **Fig**. 3c, right). Discrete annotations also allow us to identify boundaries between single-cell subcompartments and assess the transcriptional activity at regions near subcompartment boundaries. We discovered that regions at subcompartment boundaries **(Supplementary Note**) are also associated with higher transcriptional variability (**Fig**. 3c, left). SCGHOST unveils single-cell subcompartment boundaries and structurally variable chromatin regions that are both associated with high transcriptional variability.

Together, these results of applying SCGHOST to WTC11 cells further exemplify the effectiveness of our method in revealing the variability of single-cell subcompartments that are also functionally relevant.

### SCGHOST reveals single-cell subcompartments from 3D genome imaging data

We next sought to demonstrate that SCGHOST can also effectively identify single-cell subcompartments from 3D genome imaging data. We employed SCGHOST on the single-cell contact maps of ch21 at 100kb resolution, derived from chromatin tracing imaging data from MERFISH in IMR90 cells [27]. Notably, in addition to imaging the genomic loci, Su et al. [27] simultaneously imaged nascent RNA transcripts of over 1000 genes in the same individual cells, allowing us to directly compare our subcompartment annotations with gene transcription. Here, we only applied SCGHOST to the imaging data from chr21 as it is the only chromosome in the dataset that includes co-assayed single-cell gene transcription activity.

Firstly, we transformed the chromatin tracing imaging data into proximity maps, analogous to HiC contact maps, by calculating Euclidean distance maps for each cell and inverting the distance map (**Supplementary Note**). We then obtained clusters of genomic segments by using observed-over-expected proximity maps as inputs. Clusters were then matched to IMR90 population Hi-C subcompartments defined in SNIPER [3] and annotated as single-cell subcompartments. Because we only used one chromosome, we defined subcompartments using an alternative clustering method (**Supplementary Note**).

We found that the aggregated chr21 single-cell subcompartments from SCGHOST correspond to the expected enrichment of epigenomic features, further confirming the reliability of SCGHOST (**Fig**. 3d). In particular, the epigenomic enrichment is consistent with what was observed in our earlier work SNIPER [3]. In **Fig**. 3d, we observed that scA1 and scA2 are largely correlated with active histone modification and are associated with earlier replication timing. scB1, scB2, and scB3 have less active histone modification and later replication timing, with scB1 having more enrichment of H3K27me3 and earlier replication compared to scB2 and scB3. This result strongly suggests that SCGHOST reliably annotates subcompartments from 3D genome imaging data.

Next, we compared the subcompartment annotations with the number of co-assayed actively transcribed genes in each single-cell. We found that scA1 and scA2 annotations co-occur with more frequently transcribing genes **(Fig**. 3**e**, top). Moreover, scB1, scB2, and scB3 co-occur with less frequent gene transcription **(Fig**. 3**e**, bottom), suggesting that SCGHOST correctly identifies single-cell B subcompartments in population-level A compartment regions. This demonstrates that SCGHOST can identify subcompartment annotations in individual cells that directly reflect their transcriptional activity in the same cell, further elucidating the connection between large-scale chromatin spatial position and gene transcription.

Although SCGHOST is primarily intended for use on scHi-C contact maps, these results show that the method can also annotate subcompartments for single-cell 3D genome imaging data, demonstrating its broad applicability. Additionally, in the imaging dataset, single-cell subcompartment comparisons to bulk epigenomic marks remain consistent with observations made from bulk Hi-C subcompartments. Together, these results strongly suggest the reliability of SCGHOST annotations in single cells.

### SCGHOST identifies cell type-specific subcompartments in the human prefrontal cortex

A key challenge of 3D genome analysis is the identification of cell type-specific genome structure and function relationships in complex tissues. We then sought to demonstrate that SCGHOST can annotate subcompartments from scHi-C data in tissues containing a variety of cell types. Specifically, we applied SCGHOST to a scHi-C data from the human prefrontal cortex (PFC) [14] and assessed if subcompartments exhibit cell type specificity and if these cell type-specific annotations correspond with cell type-specific gene expression in this brain region.

Firstly, we evaluated SCGHOST’s ability to capture cell type-specific Hi-C contact patterns in the PFC dataset. We used the single-cell genome-wide subcompartment annotations from SCGHOST as embeddings for each cell (termed “SCGHOST embeddings”). SCGHOST improves the separation of prefrontal cortex cell types compared to Higashi scA/B compartments. We plotted the UMAP visualization of SCGHOST embeddings and Higashi scAB scores across all PFC cells (**Fig**. 4**a)**. In UMAP visualizations, we found that inhibitory neurons (Vip, Sst, Pvalb, Ndnf) and excitatory neurons (L2/3, L4, L5, L6) tend to cluster together when using Higashi scA/B, whereas with the SCGHOST subcompartments, inhibitory neurons are clearly separated from excitatory neurons. This improvement is especially noticeable for inhibitory neurons. Additionally, single-cell subcompartments within the same cell type are more similar compared to subcompartments among cells in different cell types **(Fig**. S5), further demonstrating SCGHOST’s ability to identify cell type-specific subcompartments.

**Figure 4:**
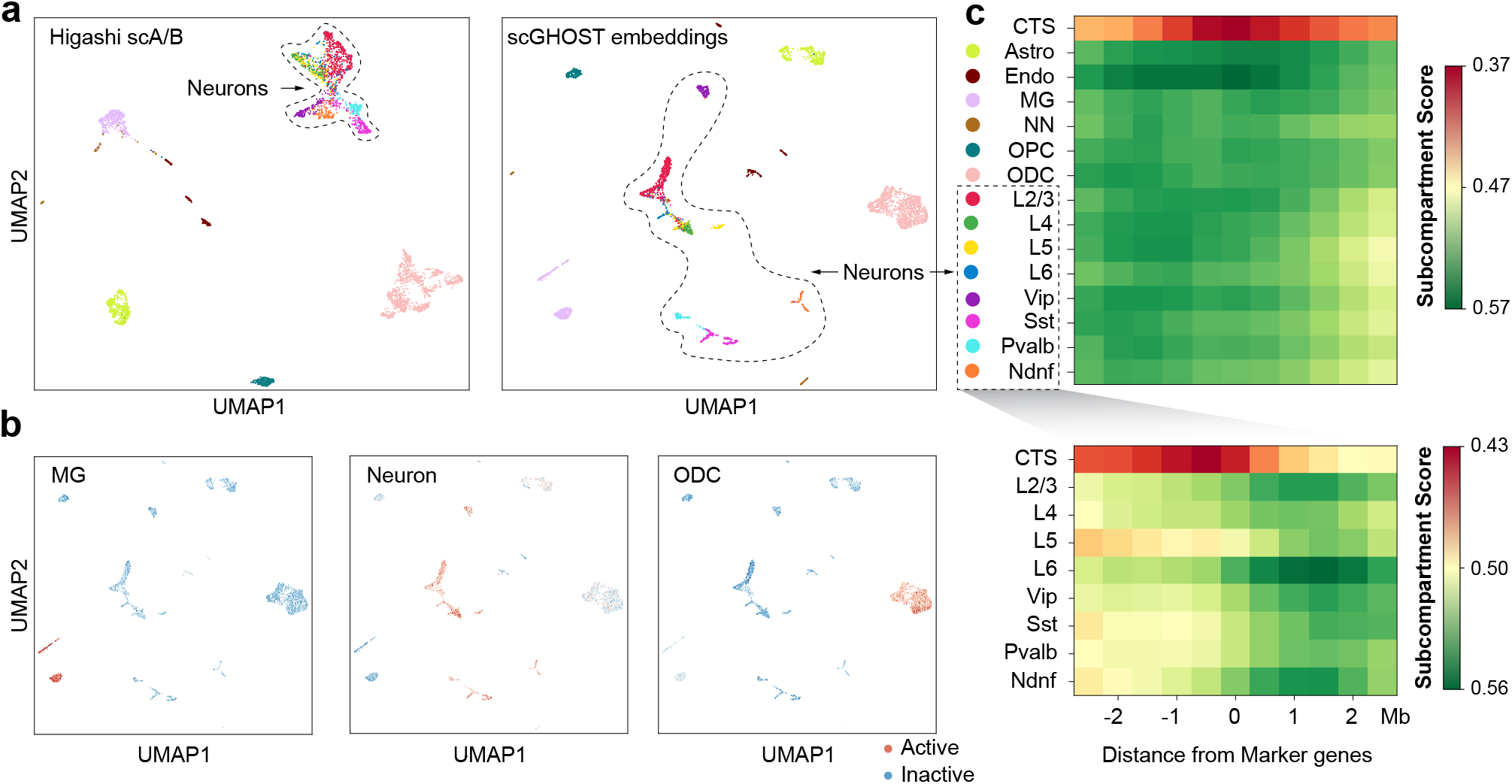
Application to scHi-C data from the human prefrontal cortex (PFC). **a**. SCGHOST more effectively separates prefrontal cortex subtypes, particularly neuronal subtypes, compared to the embeddings from Higashi scAB annotations. **b**. Subcompartment scores for the marker genes in microglia, neuron, and oligodendrocyte cells demonstrate that active subcompartments in a specific PFC subtype strongly connect with cell type-specific marker genes and can differentiate cell types. **c**. Cell type-specific (CTS) subcompartment annotations flanking marker genes in all PFC subtypes show differences in cell type-specific subcompartments that are consistent with the marker gene.

Additionally, when using a random forest classifier to directly classify PFC cell types, SCGHOST achieves accuracy comparable to using full Higashi embeddings of each cell (**Fig**. S6). However, a unique advantage of SCGHOST annotations is their direct correspondence to the genome loci. Therefore, applying classifiers that calculate feature importance (e.g., random forest) to predict cell types using SCGHOST annotations can reveal the specific genomic loci that differentiate cell types (see **Fig**. S7 for an example between L2/3 and L4). Together, SCGHOST accurately assigns single-cell Hi-C contact maps based on subcompartment patterns into different cell types, paving the way for a more interpretable understanding of single-cell 3D genome structure features, such as subcompartments, that determine cellular function.

To show that more active single-cell subcompartments in the PFC dataset contain cell type-specific marker genes, we assigned scores from 0 to 4 to five subcompartments in the PFC. Active subcompartments were assigned lower scores, while inactive subcompartments were assigned higher scores. We then gathered the subcompartment scores for the 500 marker genes (see **Methods** on the identification of marker genes) with the highest fold change in each cell type. The scatter plots in **Fig**. 4b depict the average single-cell subcompartment scores for the marker genes of specific cell types. We found that in microglia, neurons, and oligodendrocytes, their corresponding clusters in the scatter plot generally have lower scores compared to other cell types, indicating that marker genes specific to each cell type tend to locate in more active subcompartments. This analysis shows that the more active single-cell subcompartments identified by SCGHOST are uniquely associated with more active transcription of cell type-specific marker genes.

Next, for each cell type, we trained a random forest classifier to differentiate the cell type from the remainder of the population. From each trained random forest model, we located 250 genomic loci with the highest feature importance. We then found marker genes specific to each PFC cell type that also co-localized with the most important loci and compiled the subcompartments across all cells in the population within 2.5 Mb flanking regions of each marker gene. Similar to **Fig**. 4b, we assigned subcompartment scores to the flanking regions of marker genes of each cell type across all cells: active subcompartments were assigned lower scores while inactive subcompartments were assigned higher scores. For each cell type, we then calculated the average subcompartment score of each flanking region. We found that for each cell type, cell type-specific marker genes are situated in genomic loci that are significantly more active than those same loci in other cell types **(Fig**. 4c, top) with p < 0.001. This trend remains true in neuron cell types **(Fig**. 4c, bottom) that are typically more challenging to differentiate using subcompartment annotations.

In conclusion, SCGHOST effectively annotates single-cell subcompartments from scHi-C data of complex tissues. The cell type-specific subcompartments that are identified delineate different cell types, connect to cell type-specific gene regulation, and can be used to reveal specific genomic loci that modulate cellular function.

## Discussion

In this work, we developed a new method, SCGHOST, to identify single-cell genome subcompartments using scHi-C data. SCGHOST is the first computational method that provides genome-wide annotation of subcompartments in each individual cell using scHi-C data. Our extensive evaluation demonstrates that SCGHOST effectively annotates subcompartments based on scHi-C data for different cell lines, providing results consistent with functional genomic signals collected in a population of cells while also offering unique, cell-to-cell variability of subcompartments. Applications of SCGHOST to a scHi-C dataset of the human prefrontal cortex have demonstrated its ability to reveal cell type-specific subcompartments and specific chromosomal coordinates that have distinct connections to cell type-specific gene expression. In addition, an integrative analysis with data based on simultaneous imaging of chromatin architecture and the transcriptome further demonstrates that the connection between subcompartments and gene transcription exists within the same cell type at a single-cell resolution. SCGHOST uniquely captures such structure-function connections, providing a fresh perspective of the roles of subcompartment in gene regulation. Overall, SCGHOST offers a valuable new approach to enhance the analysis of single-cell 3D genome features and their implications in genome functions for a wide range of biological contexts.

There are a few possible improvements for SCGHOST. First, the framework could be improved to generate embeddings that are directly comparable across chromosomes, reducing the need for a workaround such as our approximation of single-cell inter-chromosomal Hi-C maps. Second, we can include other types of single-cell epigenomic features and functional genomic data into the model to further improve the analysis of cell type-specific characteristics of 3D genome structure and function relationships. This is particularly important to chart a more comprehensive nuclear structure-function landscape in complex tissues. Third, the discrepancy between single-cell and population-level subcompartments should be further investigated. While we are confident that single-cell annotations are specific to single-cell biological properties and offer improvements over population-level subcompartments, additional work needs to be conducted to understand the mechanisms underlying cell-to-cell 3D genome structure variability that can help reconcile single-cell and population-average discrepancies. Lastly, future work could explore whether similar results might be achieved with less complex neural network architectures. Nevertheless, we have demonstrated that SCGHOST is an effective new method for identifying single-cell subcompartments and has the potential to provide crucial insights into higher-order genome organization and function.

## Methods

### Constrained random walk sampling for SCGHOST graph embedding

Due to the noise present in imputed scHi-C contact maps, we aim to filter out contacts from scHi-C that are less informative regarding spatial interactions among genomic loci. To achieve this, we utilize random walk sampling in the contact maps with a constrained random walk space. Genomic loci pairs from the random walk sampling take the format of sparse, undirected, weighted graphs for each chromosome of each cell. These sparse graph representations are then fed input a graph embedding neural network to assign genomic loci embeddings, which we subsequently use for annotating subcompartments.

Given the scHi-C contact map M_*n*_ for cell *n ∈* [1, *N*] where *N* is the total number of cells, and maps 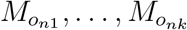 for the cells most similar to n (measured by Higashi scHi-C embeddings), we first calculate the Pearson correlation maps of the contact maps, R_*n*_, 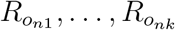 to reduce noise in the contact map. To filter out genomic loci that are less informative of spatial interactions, we next carry out constrained random walks in cell n and its neighbors at a given locus i ∈ [1, *L*] where *L* is the number of bins on a chromosome at a given resolution (**Fig**. 1b,c). For each locus, we perform *w* random walks where each random walk consists of a first and second order walk to improve the performance of the downstream graph embedding model [28]. We conduct first-order walks to sample pairs between locus *i* and its nearest neighbors, loci that share the most frequent and infrequent scHi-C contacts with *i* Second-order walks then sample second-order neighbors – loci that share the most frequent and infrequent contacts with the first-order neighbors of locus *i*. The novelty in this approach lies in selecting the first-order and second-order neighbors only selected from subsets of loci with the most informative scHiC contacts with *i*, resulting in clearer spatial connections among chromatin regions unique to different subcompartments than those offered by Higashi-imputed scHi-C maps.

To improve the performance of the subsequent graph embedding model (see **Fig**. S8 for a real data evaluation). we identify the most similar cells to each cell in the dataset (**Fig**. 1b). For each single cell *n ∈* [1, *N*], we calculate the Euclidean distances between the scHi-C embeddings of n and all other cells. The *k* cells with the lowest distance are considered the nearest cell neighbors to n. We then calculate the Pearson correlation maps of the neighboring cells, 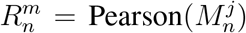, where *m ∈* [1, *k*]. Following the strategy in Higashi [17], we chose k = 5. Random walk sampling is then performed on all 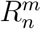 for *m ∈* [1, *k*].

#### First-order random walks

In each first-order walk, we first individually sort the rows of the Pearson correlation matrix *R*_*n*_ of cell *n* in descending order and track the indices in each row that sort the array (Fig. 1c). We compile the entries in the first *t* percentile of each sorted row and probabilistically select one genomic locus from these higher-valued entries in each row of *R*_*n*_. We then set the selected locus 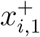 as a first-order positive sample of locus *i*. Repeat the random selection for the bottom *t* percentile, we set the selected locus 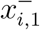 as a first-order negative sample. As a result of the first-order walk, (*i*, 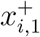) and (*i*, 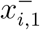) are positively and negatively connected first-order neighbor pairs, respectively.

#### Second-order random walks

SCGHOST also links loci that may not be strongly connected in the original scHi-C contact map due to noise but share strongly connected first-order neighbors, using second-order random walks. Having defined 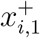 and 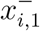, we sort the 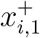 -th and 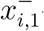 -th rows of R_*n*_ in descending order, compile the indices that sort the rows, and compile entries in the top and bottom t percentiles of those rows. We then define the second-order positive sample 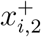 by probabilistically selecting from entries in the top t percentile of the 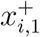 -th row and repeat this step on the 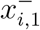 -th row to define the second-order negative sample 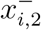. As a result of the second-order walk, (*i*, 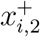) and (*i*, 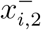) are positively and negatively connected second-order neighbor pairs, respectively.

#### Composite set of all random walks

Next, we assemble first and second-order neighbors into a format akin to sparse graphs to be inputs for a graph embedding model and subsequent subcompartment annotations (Fig. 1c). The input is represented as a set of pairs between *i* and 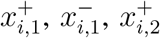, and 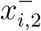 for all loci *i*. We assign labels to each pair by annotating positive pairs (*i*, 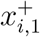) and (*i*, 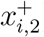) as 1 and negative pairs (*i*, 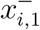) and (*i*, 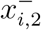) as -1. We repeat sampling of first-order and second-order walks for the Pearson correlation matrices of the *k* neighbors of cell *n* at locus *i* and define 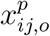 for *j ∈* [1, *k*], *p ∈* [+, *−*] denoting positive or negative samples, and *o ∈* [1, 2] denoting first or second-order neighbors of locus i. Pairs (*i*, 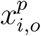) and (*i*, 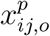), *∀ j ∈* [1, *k*], *p ∈*[+, *−*], o *∈* [1, 2] are appended to a set S containing all pairs and labels across all loci and random walks. After iterating through all loci, we filter S to only keep unique pair and label combinations.

#### Random walk parameters

For all instances of SCGHOST, we used *w* = 50 because we found that more than 50 random walks yielded little to no improvement in the training of the SCGHOST neural network. For all experiments, we set *t* to equal 25% of the number of nodes in each chromosome such that for each genomic locus, SCGHOST only performs random walks among nodes in the top and bottom 25% of contact frequencies.

### Calibrating the labels of the random walk samples

Subcompartments within the same major A/B compartment often feature similar Hi-C contact patterns but exhibit different interaction frequencies. These subtle differences are poorly captured by graph embedding if we discretely label random walk samples. Therefore, we convert discrete labels of the random walks into continuous-valued labels, allowing SCGHOST to better differentiate random walks specific to different subcompartments. We calibrate these labels by replacing the discrete label of each contact pair with its observed-over-expected Pearson correlation value. We normalize the labels by dividing positive-valued labels by the *q*-th percentile of all positive labels and dividing negative-valued labels by the 100*−q*-th percentile of all negative labels. This step is crucial to mitigate cross-chromosome bias caused by Pearson correlation distributions varying across different chromosomes. Since Pearson correlation matrices often contain values of 0 and 1, normalizing by the 100-th and 0-th percentile values would suppress the mean value of ours labels undesirably. In all instances of SCGHOST, we used *q* = 97.5, which neither over-suppresses nor over-saturates label values.

### The SCGHOST graph embedding model

We developed a graph embedding model to be used in conjunction with constrained random walks and label calibration, designed to embed noisy scHi-C contact maps (**Fig**. 1d). Random walk-based graph embedding methods [29, 30] have been applied to bulk Hi-C data in the past [23], but there has been no attempt to the scHi-C context. Our graph embedding model generates embeddings for each genomic bin (500kb in size) and aims to reconstruct cosine similarity scores between positively or negatively correlated genomic bins from these embeddings. The model consists of two neural network layers: one hidden layer between the input and the embedding output, and the embedding output layer.

The hidden layer in SCGHOST contains a *n*_*h*_ *× N*_*a*_ weight matrix 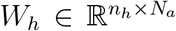 where *N*_*a*_ is the number of ungapped 500kb bins along a given chromosome a, and n_*h*_ is the dimensionality of the hidden layer. The output of the hidden layer pertaining to the i-th bin in the scHi-C contact map is 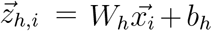, where 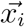 is a *N*_*a*_-dimensional one-hot encoded vector (the *i*-th entry is 1, and all other entries are 0), and *b*_*h*_ is the learnable bias term. 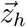 is then fed forward into the rectified linear unit activation function, which sets all negative inputs to zero.

The embedding output layer in SCGHOST contains a *n*_*out*_ *× n*_*h*_ matrix 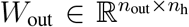 and the embedding output of the *i*-th 500kb bin 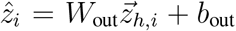, where *b*_*out*_ is the bias term of the embedding output layer and *n*_*out*_ is the output dimension.

The models are fit by inputting two different bins simultaneously, embedding both regions and computing the cosine similarity between them. For the *i*-th and *j*-th bins in the scHi-C contact map, cosine similarity between *i* and *j*, denoted by *ρ*_*ij*_, is calculated as follows:

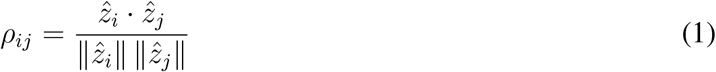

For the pair of regions *i* and *j*, the loss *l*_*ij*_ *= (ρ*_*ij*_ *− y*_*ij*_)^*2*^, where *y*_*ij*_ is the correlation between *i* and *j* as described in the scHi-C contact maps. Intuitively, the model maximizes the cosine similarity among loci that interact frequently and minimizes similarity among loci that interact infrequently. We optimize the parameters of our models using the Adam optimizer [31].

### Constructing estimated inter-chromosomal Hi-C contacts in single cells

The neural network computes embeddings independently for each chromosome. As such, clustering on a single chromosome would return clusters that might not be comparable across chromosomes. Therefore, we developed a method to approximate inter-chromosomal contacts for each single cell, facilitating cross-chromosome comparison of subcompartment annotations.

For each cell, we compute the Pearson correlation matrix of N_*a*_ 128-dimensional embeddings of each chromosome, where *N*_*a*_ is the number of genomic loci (500kb in size) in a given chromosome *a*. We then calculate the first principal component (PC1) of the correlation matrix of each chromosome, calibrate the PC1 values such that higher values correspond to the A compartment, defined using CpG density, and divide the PC1 vectors into quantiles. Rows in the upper and bottom *v*-th percentiles of each chromosome *a ∈* ***𝒞****}*, where ***𝒞*** is the set of chromosomes, are averaged column-wise into *N*_*a*_ *×* 1-dimensional vectors.

For each pair of chromosomes, we compute the outer product of the top percentiles and subtract the resulting matrix from the outer product of the bottom percentiles:

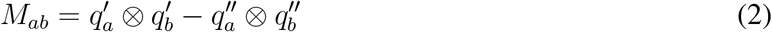

where *M*_*ab*_ is the result from the outer products between chromosomes *a* and *b* (*a, b ∈* ***𝒞***), 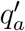 and 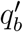 are the aggregated top percentile vectors from chromosomes a and *b*, respectively, and 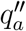 and 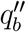 are the aggregated bottom percentile vectors. 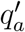 and 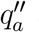 are *N*_*a*_ *×* 1-dimensional while 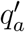 and 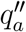 are *N*_*b*_ *×* 1-dimensional. Eq. 2 therefore returns a *N*_*a*_ *× N*_*b*_-dimensional inter-chromosomal matrix between chromosomes a and b.

*M*_*ab*_ is then quantile-normalized to have similar value ranges across different chromosome pairs. We set *a, b ∈* ***𝒞*** and concatenate *M*_*ab*_, *∀b ∈* [1, *n*_*chrom*_] horizontally:

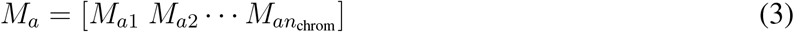

Finally, we concatenate *M*_*a*_, *∀a ∈ {*1, …, *n*_*chrom*_*}* vertically:

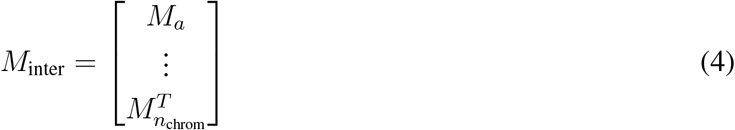

where *M*_*inter*_ is the estimated inter-chromosomal matrix.

We created the estimated inter-chromosomal contact map using odd-numbered chromosomes along the rows and even-numbered chromosomes along the columns, similar to the selections in previous annotation methods [2, 3]. Using this approach, we found that the estimated inter-chromosomal contact maps are positively correlated with population-level inter-chromosomal contact maps based on bulk Hi-C (**Fig**. S9).

### Clustering the SCGHOST embeddings

To annotate subcompartments in each cell, we begin by determining the optimal number of clusters in the genome. We applied the Bayesian Information Criterion heuristic (**Fig**. S10) on the estimated interchromosomal scHi-C contact map (see previous subsection). We used the Kneedle algorithm [32, 33] to estimate the knee point of the heuristic curve and set the horizontal axis of the knee point as the optimal cluster number.

Next, we applied Gaussian HMM clustering on the approximated inter-chromosomal scHi-C contact maps of each cell. As different subsets of the genome are present along the rows and columns of the inter-chromosomal matrices, we ran two separate clustering instances: one on the previously defined *M*_*i*_ and one on the transpose, 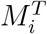 To ensure comparability of subcompartment annotations across individual cells, we sort the Gaussian HMM clustering results of each cell using PC1 values derived from Higashi.

### SCGHOST parameter selection and performance

#### Quantile selections for estimating single-cell inter-chromosomal contacts

We chose to use the upper and lower 20th percentiles to approximate the inter-chromosomal scHi-C contact map of each cell. We defined an optimal quantile as one that corresponds to (1) a high overlap between pseudo-bulk and bulk-level A and B compartments; and (2) the inflection point at which narrower quantiles offer diminishing returns in the overlap. As shown in **Fig**. S11, we found that the upper and lower 20th percentiles best satisfy these conditions.

#### Parameter selection for the graph embedding model

SCGHOST generates 128-dimensional embeddings for each genomic locus in every cell. This choice is primarily to ensure comparability with the Higashi embeddings, which also have 128 dimensions. Moreover, the SCGHOST embedding model includes a hidden layer with an output of 256 dimensions. We chose this dimensionality as it balances the input dimensionality of most chromosomes and the output dimensionality.

#### Runtime and performance

The SCGHOST framework generates a very number of sampled random walk pairs. Although each random walk is quickly generated and subsequently processed in the neural network by a GPU, the bulk of the dataset processing occurs in a single CPU thread. SCGHOST requires approximately 12 hours to run a dataset of 4,238 cells at a 500Kb resolution on a CUDA-enabled GPU and 16-core CPU. We found that the runtime of SCGHOST scales linearly with the number of cells in the dataset.

### Data acquisition and processing

In this work, we used several public scHi-C datasets. scHi-C data for the following cell lines: GM12878, HFFc6, HAP1 [12] were downloaded from the 4DN Data Portal [24, 34] (4DNES4D5MWEZ, 4DNESUE2NSGS, 4DNESIKGI39T, 4DNES1BK1RMQ and 4DNESTVIP977) in fastq format and were processed into contact maps at 500Kb resolution using the recommended processing pipeline (https://github.com/VRam142/combinatorialHiC) of the data source. The scHi-C dataset of the human prefrontal cortex [14] was downloaded from the Gene Expression Omnibus (GEO): GSE130711 in contact pairs format, which was then transformed into contact maps at 500Kb resolution. The WTC11 scHi-C dataset was downloaded from the 4DN data portal [24] (accession IDs 4DNESF829JOW and 4DNESJQ4RXY5). All scHi-C datasets were imputed with Higashi [17] (https://github.com/macompbio/Higashi) with default parameters. The imaging dataset [27] was obtained from Zenodo (https://doi.org/10.5281/zenodo.3928890). We also downloaded the scRNA-seq of multiple cortical areas of the human brain from the Allen Brain map [35, 36]. The marker genes for cell types astrocyte (Astro), oligodendrocyte (ODC), oligodendrocyte progenitor cell (OPC), endothelial cell (Endo), microglia (MG), neurons were identified using Seurat [37, 38] with the default parameters. For each cell type, the background was chosen as the rest of the cell types. When identifying marker genes for neuron subtypes, the background was chosen as the rest of the neuron cells. The genes were then ranked by the log fold-change value between a specific cell type and the background.

## Acknowledgements

This work was supported in part by the National Institutes of Health Common Fund 4D Nucleome Program grant UM1HG011593 (J.M.), National Institutes of Health Common Fund Cellular Senescence Network Program grant UG3CA268202 (J.M.), National Institutes of Health grants R01HG007352 (J.M.) and R01HG012303 (J.M.). J.M. was additionally supported by a Guggenheim Fellowship from the John Simon Guggenheim Memorial Foundation, a Google Research Collabs Award, and a SingleCell Biology Data Insights award from the Chan Zuckerberg Initiative.

## Author Contributions

Conceptualization, K.X. and J.M.; Software, K.X.; Investigation, K.X., R.Z., and J.M.; Writing, K.X., R.Z., and J.M.; Funding Acquisition, J.M.

## Code Availability

The source code of SCGHOST can be accessed at: https://github.com/ma-compbio/scGHOST.

