## Supplemental Information for "scGHOST: Identifying single-cell 3D genome subcompartments"

### Supplementary Note

#### Calculating the information content of chromatin regions

In this section, we describe how we calculated the information content of a 500kb genomic locus using subcompartment annotations of the region across all cells in a scHi-C dataset. In our work, the information content is made up of the information content of five subcompartments. At a given locus $i$, for each subcompartment, the information content is defined as:

$$I_i(s) = -p_i(s) \log \frac{p_i(s)}{p(s)} \tag{5}$$

where $I_i(s)$ is the information content of subcompartment $s$ at locus $i$, $p_i(s)$ is the proportion of subcompartment $s$ at locus $i$, and $p(s)$ is the fraction of subcompartment $s$ in the chromosome that locus $i$ belongs to.

For any given locus $i$, the aggregate information content $I_i$ is defined as the sum of information content across all subcompartments:

$$I_i = \sum_{s=1}^{k} I_i(s) \tag{6}$$

where $k$ is the number of subcompartments.

#### Calculating the *p*-values between intra- and inter-cluster contacts

For each cell in a dataset, we compute the observed-over-expected (O/E) contact map given the Higashi-imputed Hi-C contact maps of the dataset. We then iterate through each genomic locus of each single cell. For each interaction with a locus, if the interaction is between two regions in the same subcompartment, we append the O/E interaction frequency to a set of intra-cluster interactions for that subcompartment. If the interaction is between regions in different subcompartments, we append the O/E interaction frequency to a set of inter-cluster interactions for that subcompartment pair. We repeat this process for each locus and cell in the dataset.

To calculate the *p*-values between intra- and inter-cluster contacts, for each subcompartment, we use the `ttest_ind` function from the `scipy` Python package. As inputs to the function, we use the set of intra-cluster contacts for that subcompartment and the sets of inter-cluster contacts between that subcompartment and all other subcompartments. All calculated *p*-values were less than 0.001, where most calculated *p*-values exceeded floating-point precision and returned as 0.0.

### Calculating A and B compartment scores using Higashi-imputed Hi-C maps

Here, we describe how we calculated continuous A and B compartment scores from Higashi-imputed single-cell Hi-C maps. We follow the procedure as described in the Higashi paper [1]. For a given population of scHi-C maps, we construct pseudo-bulk Hi-C maps of each chromosome by summing the contacts in all single-cell maps. We then use singular value decomposition to compute a projection matrix $V^T$ of the pseudo-bulk matrix.

We then calculate the Pearson correlation matrix $R^c$ of the pseudo-bulk matrix of each chromosome $c$. We define a mean vector $\bar{R}^c$ where each element corresponds to the mean values of each column in $R^c$.

$$\bar{R}^c = \text{mean}(R^c, \text{dim=0}) \tag{7}$$

We then calculate the Pearson correlation matrices of each imputed single-cell map $\{R_1^c, R_2^c, ..., R_N^c\}$ where $N$ is the number of cells. To center the columns of each single-cell Pearson matrix, we subtract the pseudo-bulk mean vector from the columns of each single-cell matrix and compute the dot product between the resulting vectors and $V^T$:

$$\vec{P}_i^c = (R_i - \bar{R}) \cdot V^T \tag{8}$$

where $\vec{P}_i^c$ is the PC1 vector corresponding to the Pearson correlation matrix of chromosome $c$ in cell $i$. We then concatenate the PC1 vectors of each chromosome and quantile transform the resulting longer vector so that PC1 values are between 0 and 1.

### Defining variable and stable subcompartment states

We define variable and stable chromatin regions by partitioning the genome into halves where one half has lower information content (variable regions) and the other half has higher information content (stable regions), respectively. We first determine the information content of all chromatin regions in separate chromosomes. As stated in the Results, we designate chromatin regions as variable if the information content of the region is below the 50th percentile of all information content in the chromosome. Likewise, stable regions correspond to information content above the 50th percentile.

### Pseudo-bulk subcompartment annotations

Here, we clarify how pseudo-bulk subcompartment annotations are computed from a population of single-cell subcompartment annotations. We define the subcompartment annotations of a population of $N$ single cells with in chromosome $n$ as a vector $\vec{L_i^n}$. Here, $i \in \{1, ..., N\}$ and $\vec{L_i^n} = \{s_i^n(1), s_i^n(2), ..., s_i^n(M)\}$, where $s_i^n(j)$ denotes the subcompartment annotation of cell number $i$ in chromosome $n$ at locus $j \in \{1, ..., M_n\}$ ($M_n =$ the number of loci in chromosome $n$).

We then calculate a $k \times M$ matrix for chromosome $n$, $P^n \in \mathbb{R}^{k \times M_n}$ that represents the proportion of subcompartments at each genomic locus, where $k$ is the number of subcompartments:

$$P_m^n(j) = \frac{\sum_i \delta(s_i^n(j), m)}{N} \tag{9}$$

$P^n_m(j)$ is $j$-th column of the row corresponding to annotation $m$ in $P^n$ and $\delta(s^n_i(j), m)$ evaluates to 1 when $s^n_i(j)$ is equal to annotation $m$. We then compute the information content matrix $I^n \in \mathbb{R}^{k \times M_n}$:

$$I^n_m(j) = -P^n_m(j) \log \frac{P^n_m(j)}{b^n_m} \tag{10}$$

where $b^n_m$ is the background frequency of annotation $m$ in chromosome $n$:

$$b^n_m = \frac{\sum \delta(\vec{L}^n, m)}{NM} \tag{11}$$

where $\delta(\vec{L}^n, m)$ evaluates to 1 for each entry in $\vec{L}^n$. In each chromosome, we then apply Gaussian HMM clustering on the transpose of $I^n$ to obtain a $M_n$-dimensional vector of discrete annotations. We then sort annotations based on aggregated scAB scores in each cluster.

### Assigning subcompartment boundary regions in a population of single-cells

Here, we describe how we determined which regions were associated with subcompartment boundaries. Similar to variable and stable states, we set half of the genome to be boundary-associated and the other half to be non-boundary-associated. Boundary scores of a given locus are computed by counting the number of cells in which adjacent genomic loci have different subcompartment annotations from the annotation at the locus. These numbers are then divided by the total number of cells in the dataset. The regions corresponding to the top half of these values were assigned as boundary-associated regions and the remaining regions were assigned as non-boundary-associated regions.

### Converting distance maps from 3D genome imaging data to contact frequency maps

Given a set of genome locus pairs $\vec{x}_i, \vec{x}_i \in \{1, ..., M_c\}$ and the corresponding 3D genome coordinates (from [2]) $\vec{X}_i, \vec{Y}_i \in \mathbb{R}^{N_i \times 3}$ where $N_i$ is the total number of pairs for cell $i$ and $M_c$ is the number of loci in chromosome $c$, we first compute distance maps for each chromosome $c$, denoted $\boldsymbol{\mathcal{D}}_i \in \mathbb{R}^{M_c \times M_c}$. Each entry in $\boldsymbol{\mathcal{D}}_i$ is equal to the pairwise Euclidean distance between two genomic loci:

$$\boldsymbol{\mathcal{D}}_i(\vec{x}_i, \vec{y}_i) = \|\vec{X}_i - \vec{Y}_i\| \tag{12}$$

The distance map calculation is repeated for all cells. To convert the distance map to the contact frequency map of cell $i$, denoted $\boldsymbol{\mathcal{C}}_i$, we use the inverse of the distance map:

$$\boldsymbol{\mathcal{C}}_i = \frac{1}{\boldsymbol{\mathcal{D}}_i} \tag{13}$$

### Clustering on a single chromosome for chr21 of the IMR90 imaging dataset

To annotate only one chromosome, chr21, in the 3D genome imaging dataset of IMR90rom [2], we could not approximate the inter-chromosomal Hi-C matrix as we did not have data for other chromosomes at the same resolution. Instead, we first calculate the Pearson correlation matrices $\boldsymbol{\mathcal{R}}_i$ of all chr21 scHi-C contact maps at 100kb resolution.

We then construct a stacked matrix comprised of all $\boldsymbol{\mathcal{R}}_i$ by vertically concatenating $\boldsymbol{\mathcal{R}}_i, \forall i \in \{1, ..., N\}$ to form a single $NM_c \times M_c$ matrix, where $N$ is the number of cells in the scHi-C dataset and $M_c$ is the dimensionality of each Pearson correlation matrix. We then apply Gaussian HMM clustering from the hmmlearn Python package on the stacked matrix: $\boldsymbol{\mathcal{S}} = \left[\boldsymbol{\mathcal{C}}_1^T, \cdots, \boldsymbol{\mathcal{C}}_N^T\right]^T$, where $\boldsymbol{\mathcal{C}}_i$ is the contact frequency map of cell $i$ in the imaging dataset.

The result is a set of subcompartment annotations for each cell that does not need to be re-aligned to be comparable across the population. However, each cluster from Gaussian HMM does not necessarily correspond correctly to active and inactive subcompartments. As such, we then sort the annotation set based on the mean first principal component value of each single-cell subcompartment.

To compute the first principal component of each subcompartment, we first calculate the projection matrix of the population Hi-C chr21 matrix (see previous subsection). For each cell, we then compute the dot product between the correlation matrix of each cell and the population-level projection matrix and bin each PC1 value to their associated subcompartment annotations. For each subcompartment, we can then determine the mean PC1 value and use the argsort method in numpy to determine the order of single-cell subcompartments from most active to least active.

### Mean population distribution of stable and variable subcompartments

Here we explain how we determined the distribution of subcompartments of genomic regions labeled as stable or variable in a single-cell dataset. Given sets of stable and variable chromatin regions:

$$X_{\text{stab}} = \begin{bmatrix} \text{chr}_1 & \text{start}_1 & \text{end}_1 \\ \text{chr}_2 & \text{start}_2 & \text{end}_2 \\ & \cdots & \\ \text{chr}_{N_{\text{stab}}} & \text{start}_{N_{\text{stab}}} & \text{end}_{N_{\text{stab}}} \end{bmatrix}; X_{\text{var}} = \begin{bmatrix} \text{chr}_1 & \text{start}_1 & \text{end}_1 \\ \text{chr}_2 & \text{start}_2 & \text{end}_2 \\ & \cdots & \\ \text{chr}_{N_{\text{var}}} & \text{start}_{N_{\text{var}}} & \text{end}_{N_{\text{var}}} \end{bmatrix}$$

where $N_{\text{stab}}$ and $N_{\text{var}}$ are the number of loci in stable and variable states, respectively. The start and end entries define the interval in the chromosome that is either stable or variable. We iterate over both sets of chromatin intervals and calculate the overlap of stable and variable states with population subcompartments.

We compile the total number of 100kb population subcompartment regions that lie within stable or variable states and calculate the frequency of each subcompartment within each state:

$$f_{\text{stab}} = \frac{\begin{bmatrix} N_{\text{stab}}^{A1} & N_{\text{stab}}^{A2} & N_{\text{stab}}^{B1} & N_{\text{stab}}^{B2} & N_{\text{stab}}^{B3} \end{bmatrix}}{\sum N_{\text{stab}}^{s}} \tag{14}$$

where $N_{\text{stab}}^{s}$ refers to the number of 100kb loci in each subcompartment $s \in \{\text{A1, A2, B1, B2, B3}\}$ in stable in the cell population. We used the same calculation for frequency in variable states, $f_{\text{var}}$. To compute the values in **Fig.** S3a, we compute the genome-wide frequency of each subcompartment then divide $f_{\text{stab}}$ and $f_{\text{var}}$ by the genome-wide frequency.

#### Calculating marker gene enrichment in human PFC cells

Here we explain how we computed the marker gene enrichment shown in **Fig.** 4b. Each cell in the population has a discrete annotation set given as $L_i$ for $i \in [1, N]$, where $N$ is the number of cells in the single-cell dataset. Certain cell types such as neuron cell types have proportionally more genomic regions in the active subcompartments than other cell types (**Fig.** S12). Therefore, we normalize the bias in these cell types by quantile transforming the discrete annotations of each cell. As a result, $L_i$ in each cell has a value range between 0 and 1.

For each cell type, we then compile the chromosomal locations of up to 500 marker genes with the largest fold change over control. At the 500kb loci of each marker gene, we compile the subcompartment annotations for all cells. After iterating through all marker genes of a cell type, we compute the average subcompartment label of each cell and partition the cells by cell type. We then plot the UMAP of the scGHOST embeddings and color code each cell based on the average subcompartment label of that cell. Cells corresponding to red dots on the UMAP (**Fig**. 4b) indicate a more active mean subcompartment label whereas blue dots indicate a cell with inactive mean subcompartment labels. See "Data acquisition and processing" section in the main text on how marker genes were identified.

### Supplemental Figures

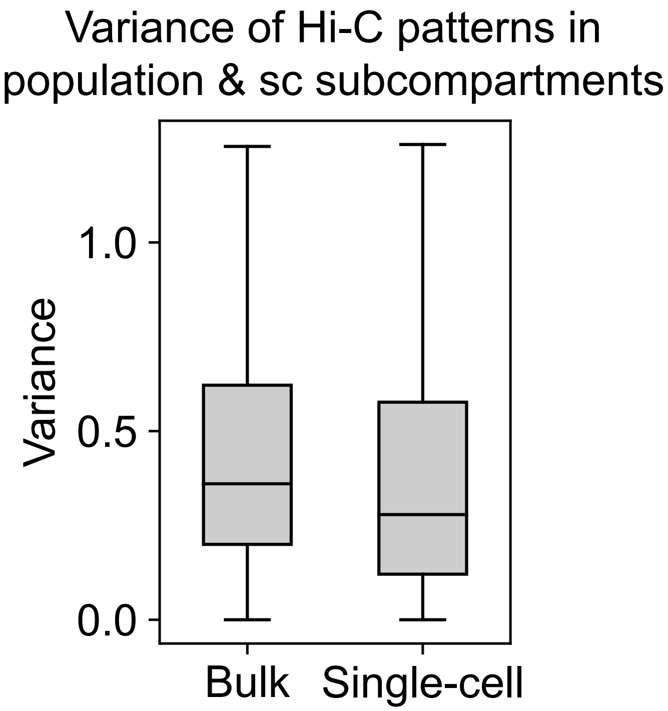


**Figure S1:** Pseudo-bulk Hi-C contact patterns in aggregated single-cell subcompartments (right) have lower variance compared to subcompartment annotations from bulk Hi-C (left) with one-sided $p < 0.001$.

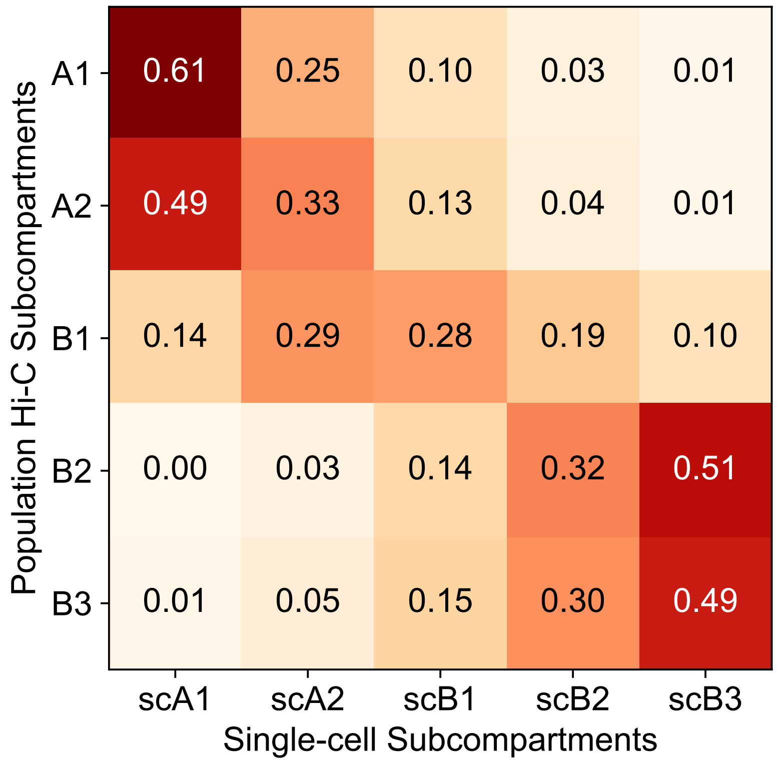


**Figure S2:** Confusion matrix between bulk Hi-C and scHi-C subcompartment annotations. The level of overlap is measured by the number of single-cell subcompartment annotations in the population of cells that share the same genomic region as a bulk Hi-C subcompartment annotation. We then divide the number of annotations for each single-cell subcompartment by the total number of annotations.

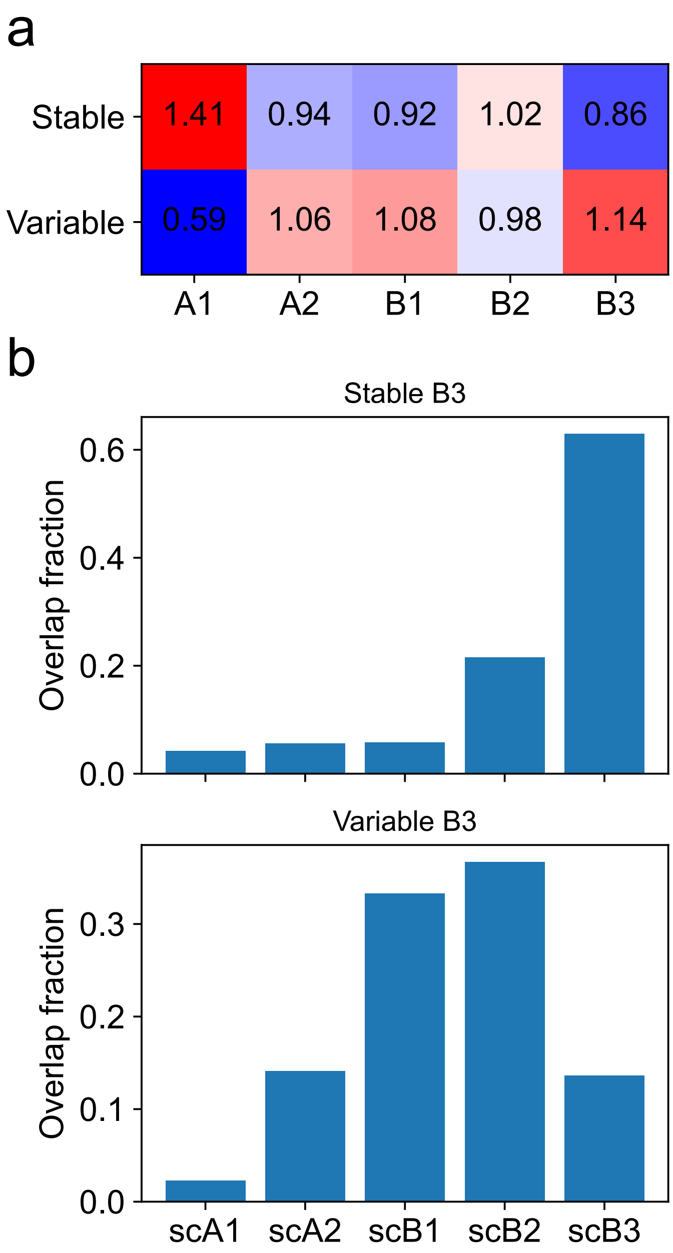


**Figure S3:** Stable and variable single-cell subcompartments that overlap with different population-level subcompartments. **a**. Mean enrichment of bulk Hi-C subcompartments in genomic regions with stable and variable single-cell subcompartments. **b**. Distribution of single-cell subcompartments in stable B3 (based on bulk Hi-C annotation; top) and variable B3 (bottom).

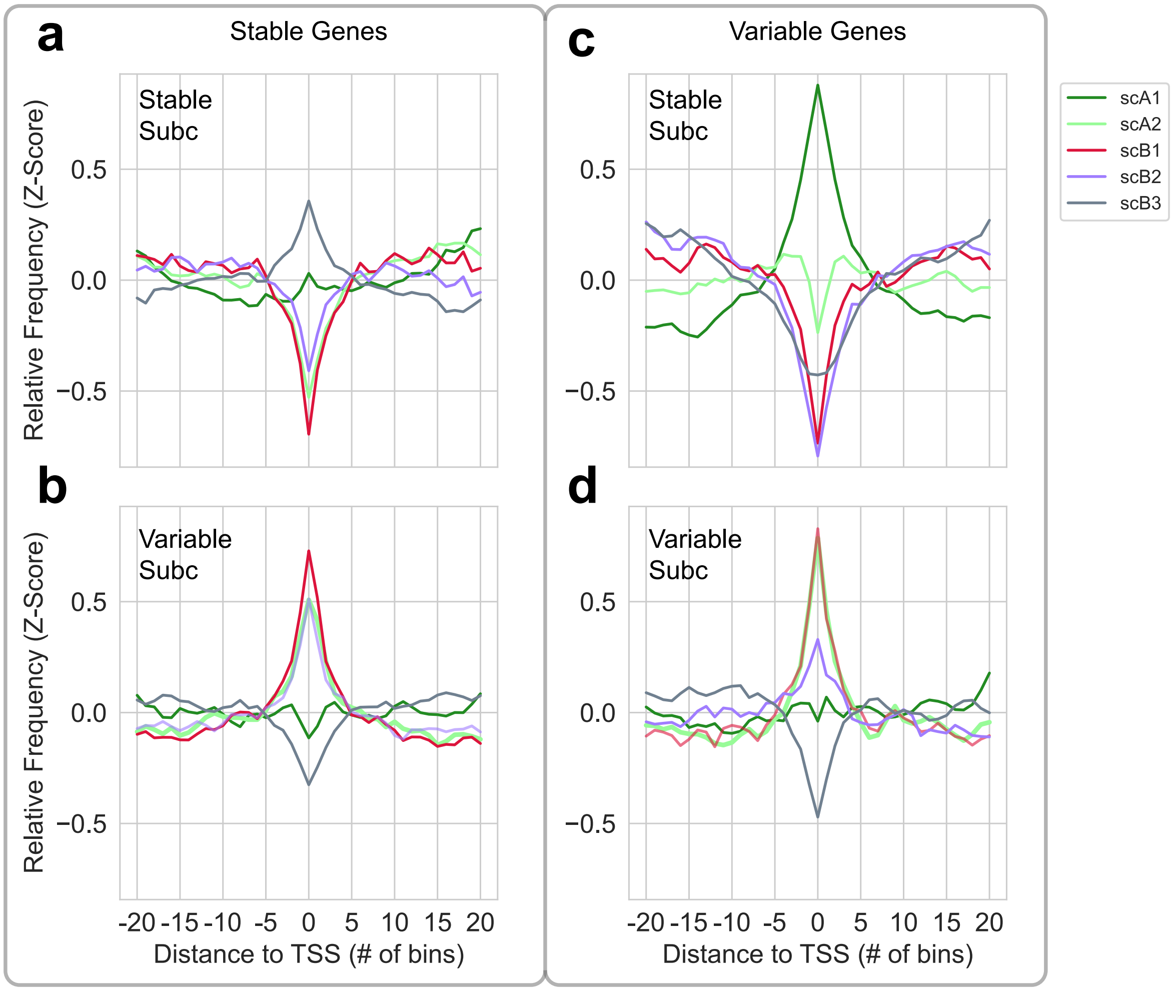


**Figure S4:** Stably (left) and variably (right) transcribed genes are associated with different single-cell subcompartments in genomic loci with stable (top) and variable (bottom) subcompartments. **a**. Stably transcribed genes in stable subcompartments are more frequently in the scB3 subcompartment. **b**. Stably transcribed genes in variable subcompartments are more frequently in scA2, scB1, and scB2 with less presence in scB3. **c**. Variably transcribed genes in stable subcompartments are more frequently in scA1. **d**. Variably transcribed genes in variable subcompartments are more frequently in scA2 and scB1.

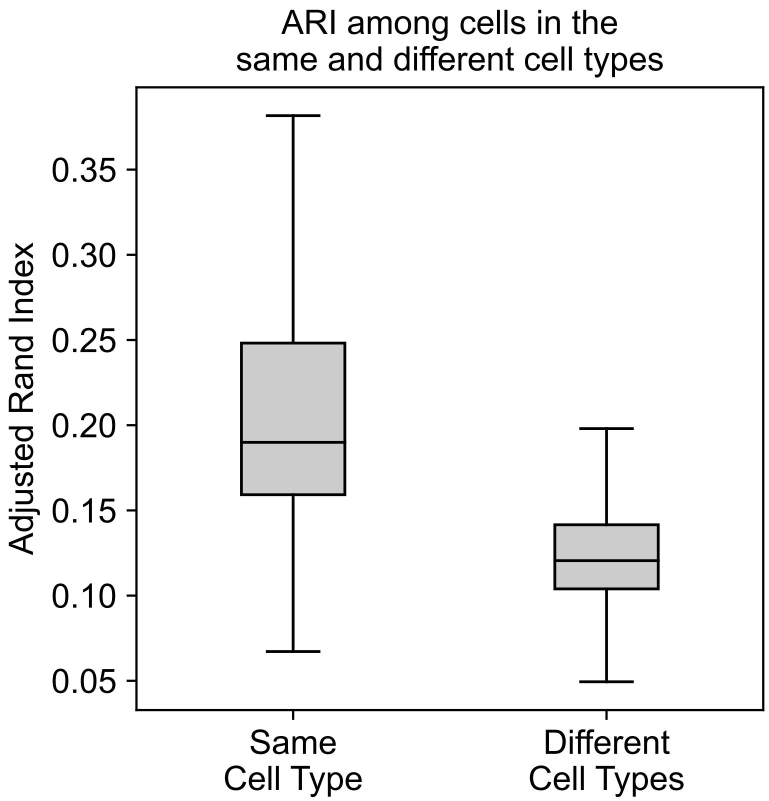


**Figure S5:** Adjusted Rand Index scores among single cells in the same cell type (left) from the human prefrontal cortex (PFC) and among cells in different cell types (right).

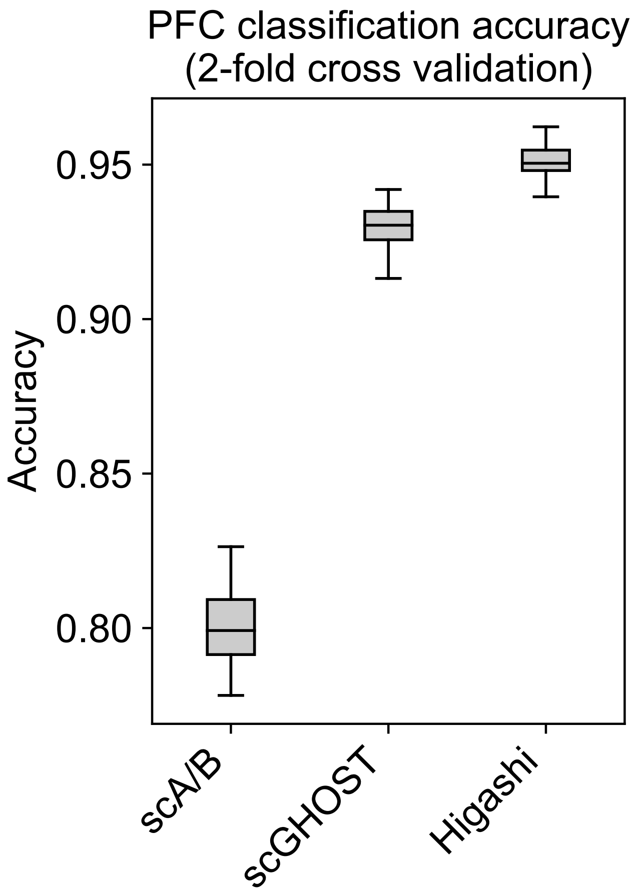


**Figure S6:** Prediction accuracy of the human prefrontal cortex (PFC) cell types using Higashi scA/B scores, SCGHOST subcompartment annotations, and Higashi single-cell embeddings. We performed 50 instances of 2-fold cross validation by randomly selecting half of the cells in the population as the training set and setting the remaining cells as the validation set.

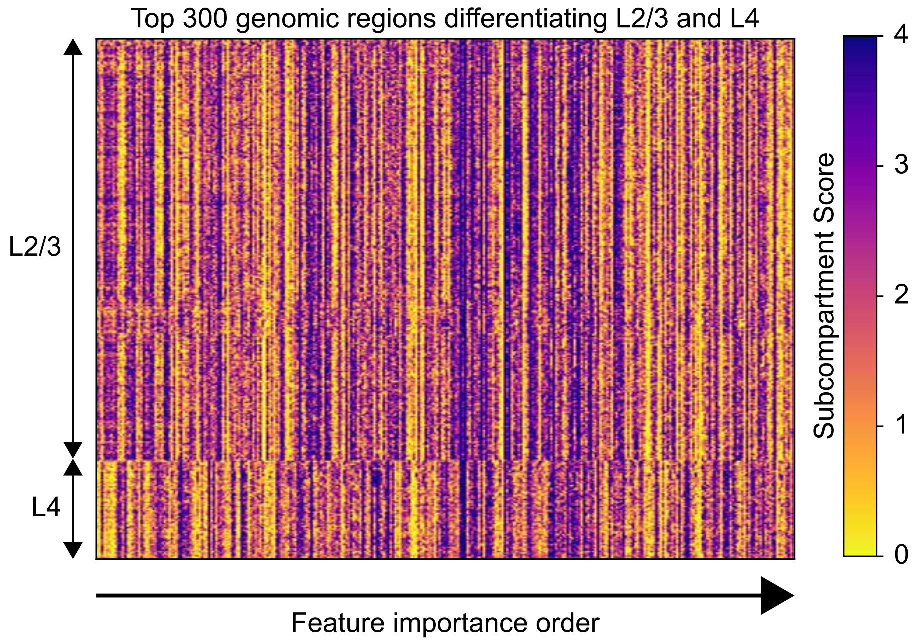


**Figure S7:** Random forest classifiers applied on the subcompartment annotations of L2/3 and L4 cells reveal the most important genomic loci that differentiate the two cell types.

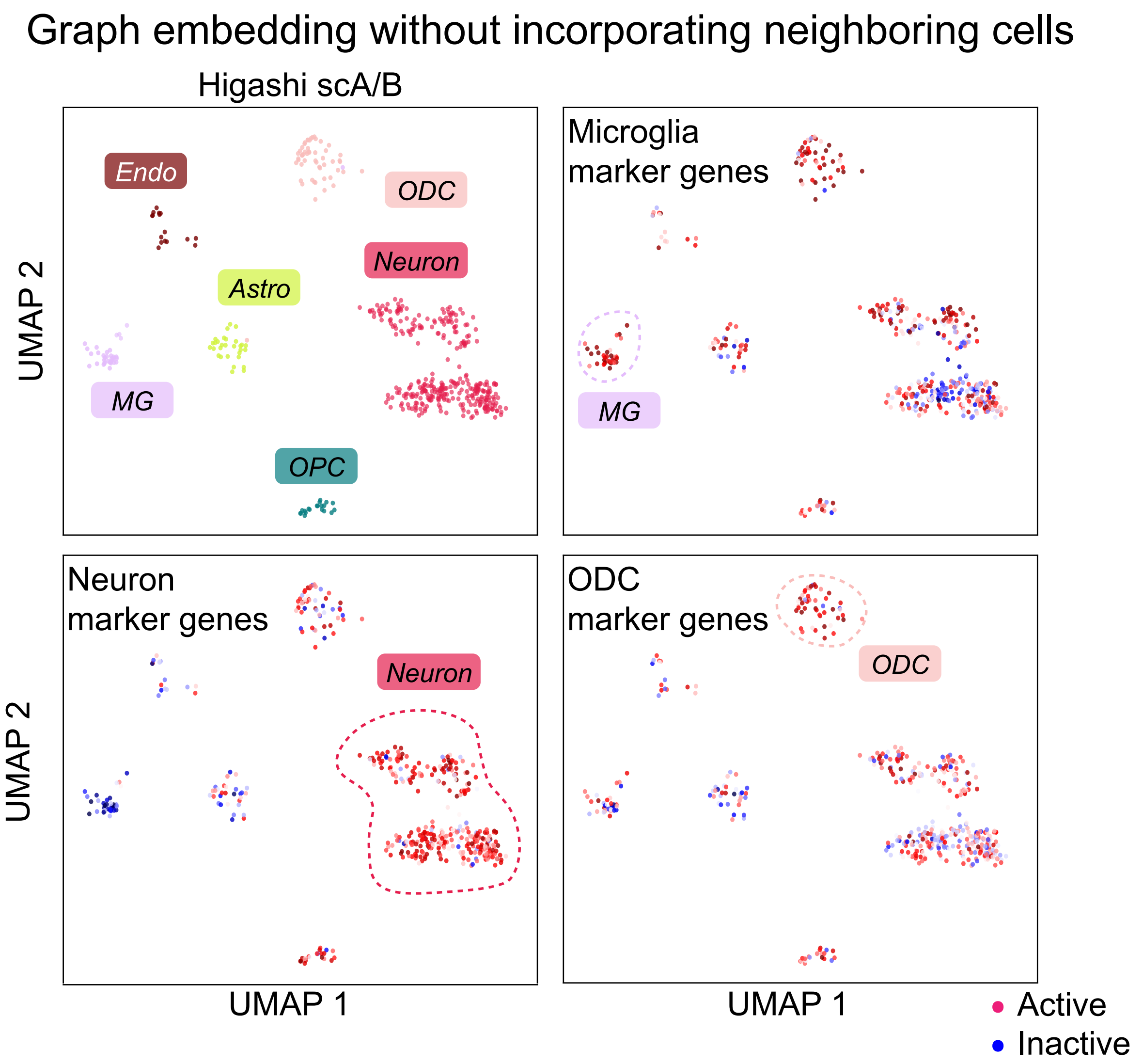


**Figure S8:** When applied to the human pre-frontal cortex, scGhost poorly separates cell types when trained without using neighboring cells. In contrast to **Fig.** 4b, here marker genes of prefrontal cortex subtypes are not in cell type-specific active subcompartments.

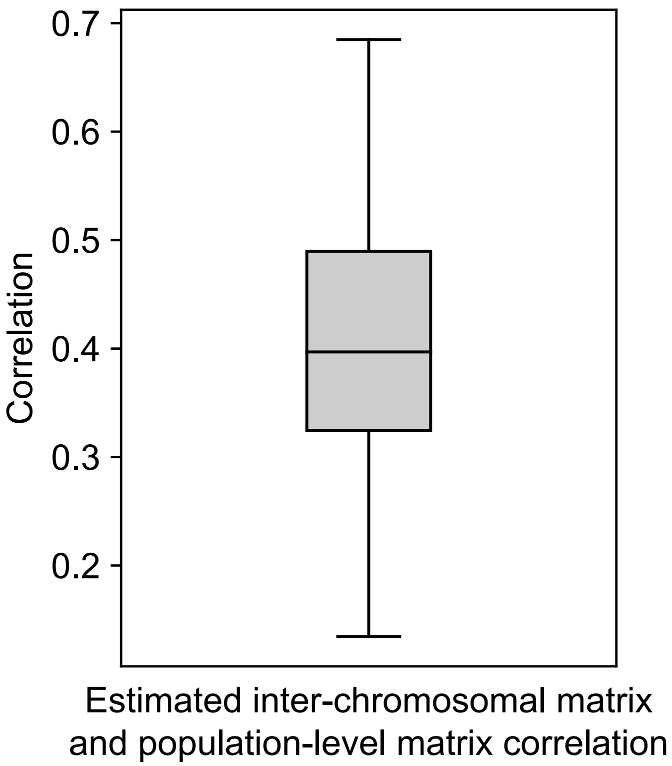


**Figure S9:** Chromosome pairs in the estimated GM12878 inter-chromosomal matrix are positively correlated with the same pairs in the population-level map.

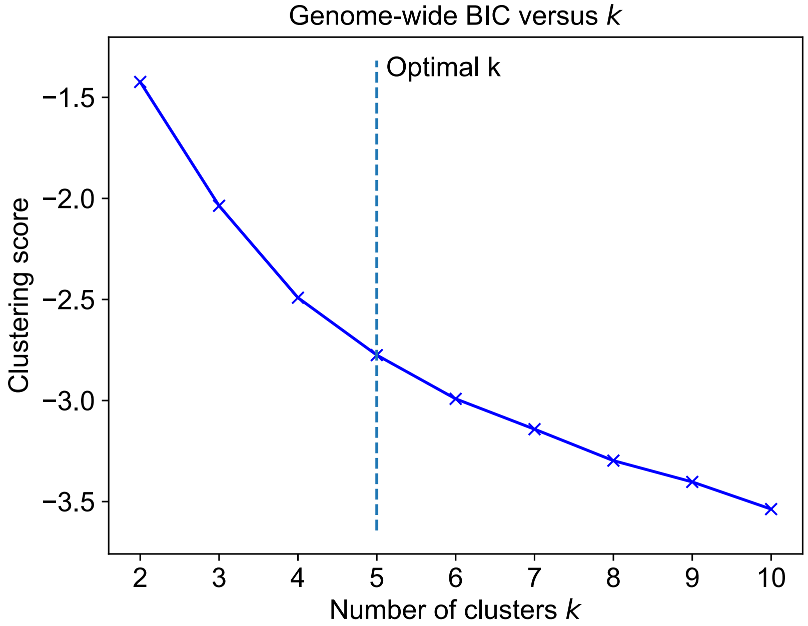


**Figure S10:** Bayesian Information Criterion (BIC) of the Gaussian HMM parameters after training on $k \in [2, 10]$ clusters. The Kneedle algorithm detected that $k = 5$ is considered as the optimal genome-wide cluster number.

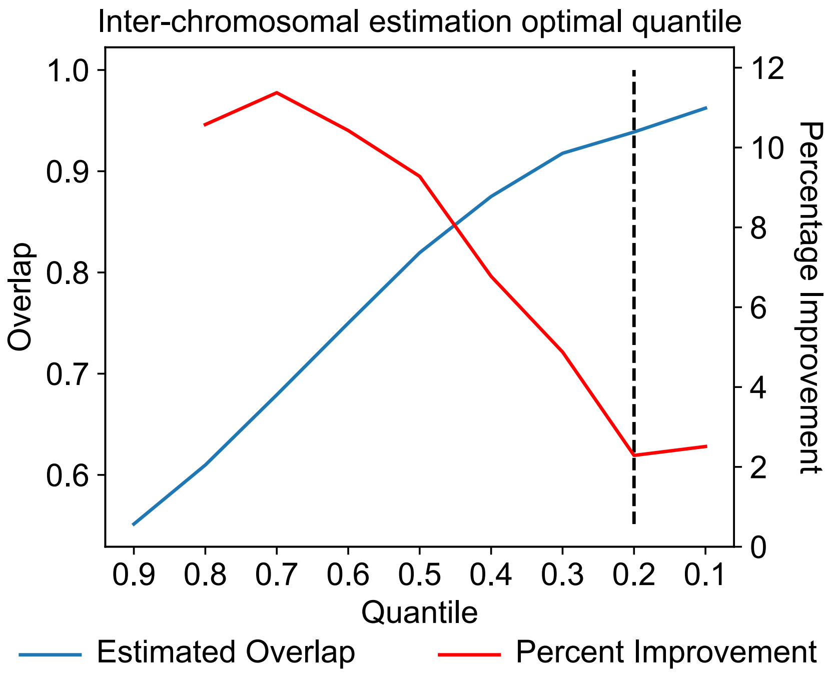


**Figure S11:** Determining the optimal quantile to use for single-cell inter-chromosomal contact estimations. The dotted line segment denotes the quantile at which: (1) the A and B compartments in estimated and bulk inter-chromosomal Hi-C have high overlap; and (2) the percentage improvement of overlap compared to the previous quantile is minimized.

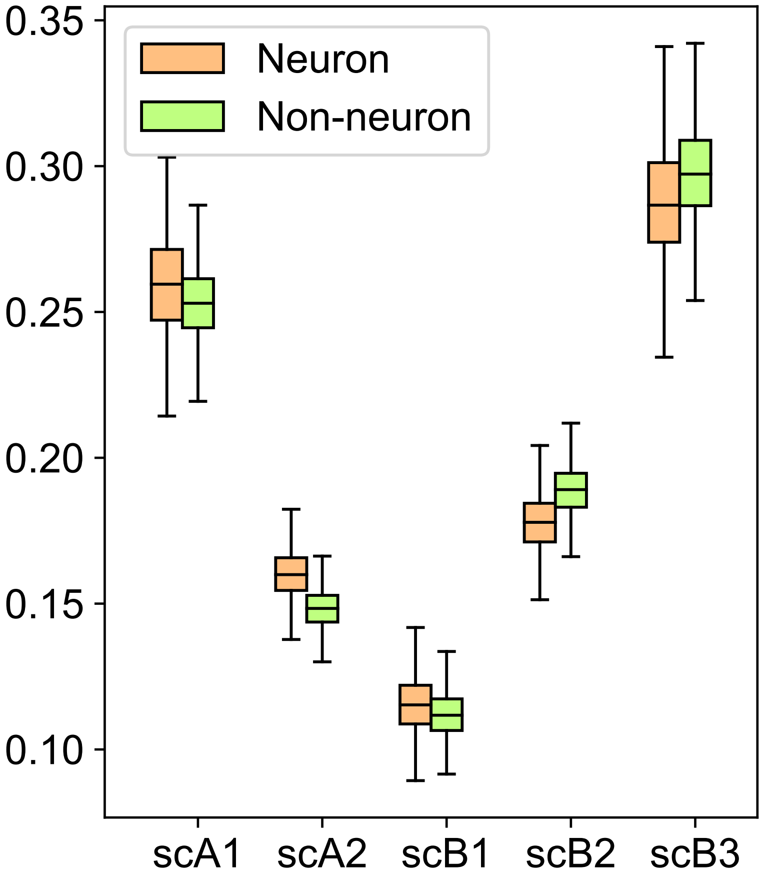


**Figure S12:** Fraction of single-cell subcompartments in neuronal and non-neuronal cell types in the human prefrontal cortex dataset. *P*-values in all subcompartments between neuronal and non-neuronal cells were measured as $<$0.001.
